# TLR3 Activation of Microglia-Containing Cerebral Organoid Induces Antiviral Factors against HIV-1 Infection

**DOI:** 10.64898/2025.12.17.695025

**Authors:** Qian-Hao Xiao, Xu Wang, Priyanka Sarkar, Xiao-Long Wang, Li Song, Binhua Ling, Wen-Hui Hu, Wen-Zhe Ho

## Abstract

Human induced pluripotent stem cell (iPSC)-derived cerebral organoids have been increasingly used as a brain model for studying various neurological disorders and neurotropic virus infections, including HIV-1. However, it is unclear whether iPSC-derived cerebral organoids possess functional innate antiviral immunity against HIV-1. In this study, we examined Toll-Like Receptor 3 (TLR3) activation and its role in the induction of IFNs and IFN-stimulated genes (ISGs) against HIV-1 in human iPSC-derived microglia containing cerebral organoids (MCOs). We observed that MCOs possess functional TLR3, which could be effectively activated by poly I:C. TLR3 activation of MCOs resulted in HIV-1 inhibition and induction of IFN-β/IFN-λ, antiviral ISGs (MX1, MX2, GBP5, SAMHD1, Viperin, and ISG56), and CC chemokines (MIP-1α, MIP-1β, and RANTES), the ligands for HIV entry coreceptor CCR5. This TLR3 activation-mediated anti-HIV-1 effects could be blocked by a specific TLR3 inhibitor. These findings indicate that human iPSC-derived MCOs are a suitable model for investigation of brain immunity and HIV-1 infection.

## 1. Introduction

Toll-like receptor 3 (TLR3) in conjunction with TLR7 and TLR9 constitutes an efficient sensor system to detect viral infections. While TLR7 and TLR9 can be triggered by single-stranded RNA viruses and cytosine phosphate guanine DNA, respectively (Cen et al., 2013; Guo et al., 2018; Meng et al., 2021; Xu et al., 2019), TLR3 recognizes double-stranded RNA viruses (Tissari et al., 2005) and triggers antiviral responses. TLR3 activation can be mimicked in experimental settings using synthetic analogs such as poly I:C. Importantly, TLR3 activation by its ligands triggers IFN signaling pathway and induces the production of both type I and type III IFNs as well as IFN-stimulated genes (ISGs). Although HIV is a single-strained RNA virus that can be detected by TLR7 (Meng et al., 2021), it can be recognized by TLR3 as well (Brulé et al., 2002; Y. Zhou et al., 2010) because HIV forms double-stranded RNAs during its replication. In HIV-1 infection of the brain, viral RNA released from either outside the blood brain barrier or HIV-1-infected cells can activate the TLR-IFN signaling pathway in microglia, triggering the antiviral response in the brain. A study showed that primary human fetal microglial cells possess TLR3 which could be activated and induce IFN regulatory factor 3 (IRF3)-dependent anti-HIV effect (Suh et al., 2009). We reported that TLR3 activation potently inhibited HIV infection of primary human macrophages through producing the multiple intracellular viral restriction factors (Dai et al., 2015; Sang et al., 2014; Y. Zhou et al., 2010). In addition, we recently reported that TLR3-IFN signaling activation of human iPSC-derived microglia could induce antiviral cellular factors against HIV-1 (Wang et al., 2023). While these observations with macrophages and microglia cultures are significant, further studies using a human brain model are necessary to verify the role of TLR3 in the brain innate immunity against HIV-1.

The brain innate immunity depends primarily on the functions of glial cells, especially microglia which are important for the early control of viral replication and clearance. Microglial cells produce antiviral factors against viral infections, including HIV-1. Therefore, the brain may rely heavily on innate immune response to prevent and control HIV-1 infection and establishment of HIV-1 latency (Geffin & McCarthy, 2013). In the brain microenvironment, microglial functions significantly depend on their direct and/or indirect contact with other brain cell types such as neurons and astrocytes. Therefore, it is clinically relevant and significant to develop a 3D brain organoid model with all major brain cell types. Recently, human induced pluripotent stem cell (iPSC)-derived cerebral organoids have been increasingly used as a brain model for studying various neurological disorders and neurotropic virus infections. Several groups have demonstrated that the human brain organoids with microglia are susceptible to HIV-1 infection (Donadoni et al., 2024; Dos Reis et al., 2022; Gumbs et al., 2022; Kong et al., 2024; Narasipura et al., 2025; Wei et al., 2023). However, it is unclear whether these brain organoids possess functional innate antiviral immunity against HIV-1 infection. In this study, we examined whether human iPSC-derived microglia containing cerebral organoids (MCOs) express functional TLR3 which could be activated by poly I:C and IFNs and IFN-stimulated genes (ISGs). In addition, we examined the mechanisms for TLR3 activation-driven HIV-1 inhibition in MCOs.

## 2. Materials and methods

### 2.1 Human pluripotent stem cell cultivation

Experiments were performed utilizing four different human iPSC lines: WT10, WT11, WT15, which obtained from Human Pluripotent Stem Cell Core at the Children’s Hospital of Philadelphia (CHOP) (Philadelphia, 2024), and the iPSC line IPS11 was generated from ALSTEM (Cat# iPS11, ALSTEMBIO). Prior to process inoculation, iPSCs were expanded in feeder-free monolayer culture on Matrigel coated plates (STEMCELL technology). All the iPSC lines were cultured in a 6-well plate as monolayer on 0.1mg/mL Matrigel (Corning) in DMEM/F12 (Gibco) media. The cells were maintained in mTeSR™ Plus media (STEMCELL Technologies) and kept at 37°C with 5% CO_2_ with fresh media change every other day. Once 60-70% confluency was reached, the cells were split with mechanical dissociation by using 0.5mM ultrapure EDTA (Invitrogen) in PBS.

### 2.2 Human iPSC-derived microglia containing cerebral organoids (MCOs)

Human iPSC-derived cerebral organoids were generated with the STEMdiff™ Cerebral Organoid Kit (STEMCELL technology) as depicted in Figure. 1A. On day 0, iPSC colonies were dissociated into single cells with Versene (Gibco). A total of 9000 cells/well were plated in ultra-low attachment 96-well plates (Corning Costar) and maintained in 100µL of EB Seeding Medium (STEMdiff™ Cerebral Organoid Basal Medium 1, supplemented with STEMdiff™ Cerebral Organoid Supplement A, and 10 µM Y-27632). At day 2 and 4, 100µL of EB Formation medium (STEMdiff™ Cerebral Organoid Basal Medium 1 supplemented with STEMdiff™ Cerebral Organoid Supplement A) was added to the culture. At day 5, embryoid bodies were transferred to a 24-well ultra-low attachment plate (Corning Costar) and cultured in Induction Medium (STEMdiff™ Cerebral Organoid Basal Medium 1 and STEMdiff™ Cerebral Organoid Supplement B). At day 7, embryoid bodies were transferred into cold Matrigel (hESC-qualified Matrix, Growth Factor Reduced, BD Matrigel™, Corning) droplets and subsequently incubated at 37°C with 5% CO_2_ for 30 min to induce Matrigel polymerization. Then, the Matrigel droplets embedding the embryoid bodies were incubated in Expansion Medium (STEMdiff™ Cerebral Organoid Basal Medium 2 with STEMdiff™ Cerebral Organoid Supplement C and STEMdiff™ Cerebral Organoid Supplement D) in a 6-well ultra-low attachment plate (Corning Costar). At day 10, organoids were placed in maturation medium (STEMdiff™ Cerebral Organoid Basal Medium 2 and STEMdiff™ Cerebral Organoid Supplement E in a 49:1 ratio) in an incubator with orbital shaker (65 rpm), at 37°C with 5% CO_2_, and kept in these conditions until the end of the experiment.

**Figure 1.**
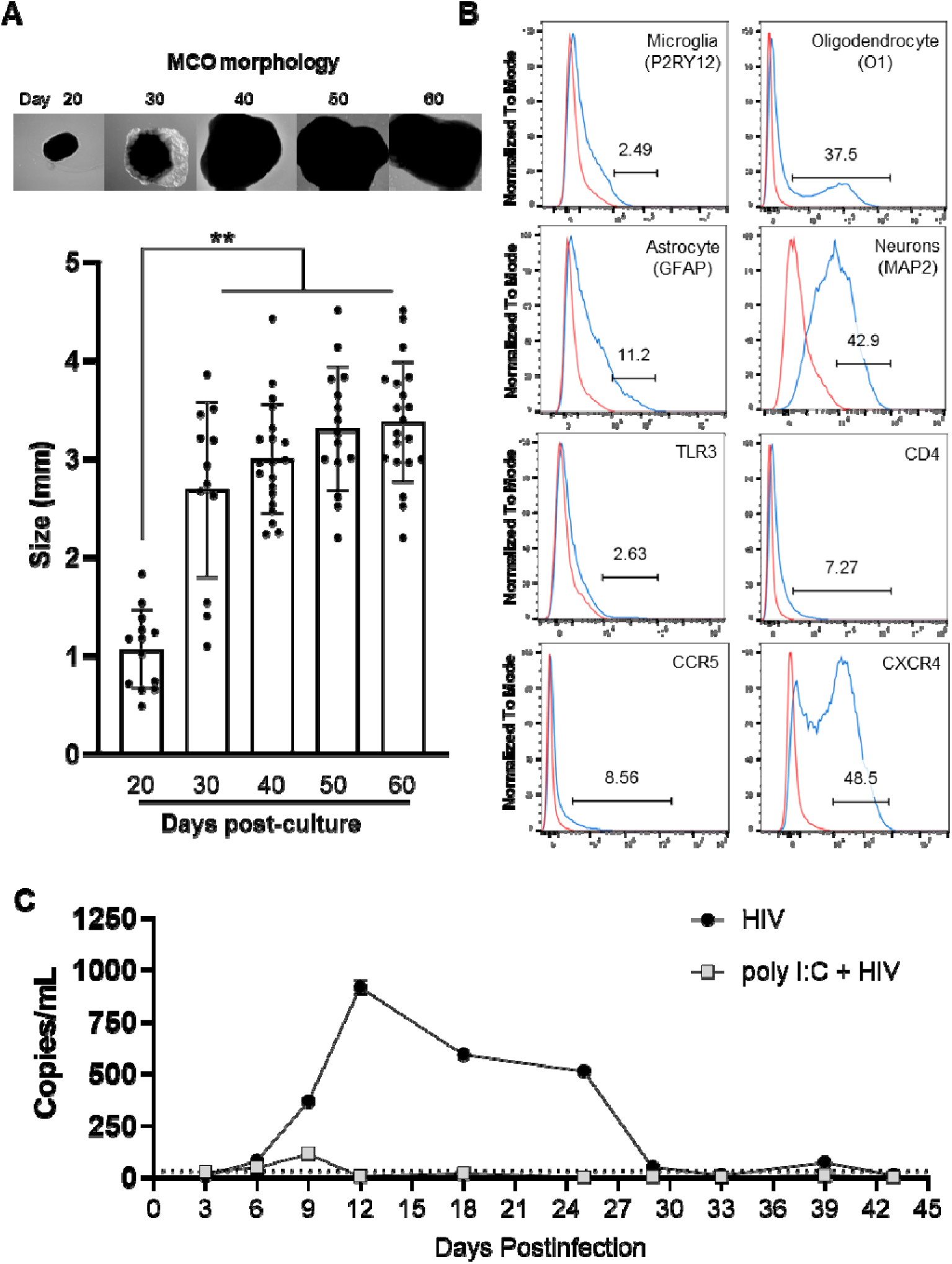
Characterization and effect of poly I:C on HIV infection of microglia-containing organoids (MCOs). (**A**) MCOs morphology from a representative iPSC line donor (upper panel) and size distribution (lower panel) during 60-day culture. Each dot represents a single organoid, and size (mm) was measured using ImageJ. Data are presented as mean ± SD relative to organoids at the age of Day 20. Asterisks indicate significant differences between groups (**P < 0.01). (**B**) Flow cytometry analysis of cell markers, HIV entry receptors and TLR3 in MCOs. As indicated: P2RY12 (microglia), O1 (oligodendrocyte), GFAP (astrocyte), and MAP2 (neuron). In addition, HIV entry receptors (CD4, CCR5, CXCR4) and TLR3 positive cells were examined. Of note: percentages of positive cells were calculated from viable singlet cells. (**C**) 50 day-cultured MCOs were pretreated for 12h with the TLR3 agonist Poly I:C (10 µg/mL), then infected with live HIV-1_Bal_ (2ng p24 per well in 48-well plate) for 48h. Cultures were maintained with poly I:C, and viral replication was monitored by measuring HIV Gag RNA in the supernatant over the indicated infection period. Data is represented as mean ± SD of triplicate cultures, representative of two independent experiments.

### 2.3. MCOs morphology

Brightfield micrographs were captured using an EVOS imaging system (Thermo Fisher Scientific) and subsequently analyzed using ImageJ (National Institutes of Health, Bethesda, MD, USA). Diameter of organoid was measured every 10 days. The size was calculated as the average of the major axis and its orthogonal. Organoids were excluded from measurements if: **i**) the center of the vesicle-like object was opaque, **ii**) debris obscured ability to define inner and outer epithelial boundaries, and/or **iii**) the structure was incomplete.

### 2.4. PolyI:C and TLR3 inhibitor treatment

The experiments were conducted with MCOs at age of 50-60 days. To activate TLR3, polyinosinic-polycytidylic acid (poly I:C, HMW, Invivogen) was mixed with LeyVoc (Invitrogen, USA) and the mixture was added into MCOs cultures. To inhibit TLR3 activation, MCOs were incubated with the TLR3/dsRNA complex inhibitor (TCI, Merck, 614310) for 4h prior to addition of poly I:C. At the indicated time points, MCOs were harvested for RNA extraction or protein analysis.

### 2.5. Cell Viability Assay

Single cells dissociated from MCOs were plated in 96-well plates (2 × 10^4^ cells/well) and treated with poly I:C or TLR3/dsRNA complex inhibitor (TCI) for 48h at 37°C. CellTiter 96® Aqueous One Solution Cell Proliferation Assay Kit 8 (Promega) was used to measure the cells viability.

### 2.6. Quantitative PCR

Total RNA from MCOs was extracted using AllPrep DNA/RNA/Protein Mini Kit according to handbook. Quantification of RNA sample concentrations and purity was measured using the NanoDrop spectrophotometer (Thermo Fisher Scientific). For each sample, total RNA (1 µg) was subjected to reverse transcription (RT) using reagents from Promega (Promega, WI, USA). The RT system with random primers for 1h at 37°C. The reaction was terminated by incubating the reaction mixture at 99°C for 5 min, and the mixture was then kept at 4°C. The resulting cDNA was then used in triplicate reactions of quantitative PCR reactions (RT-PCR). The RT-PCR was performed with iQ SYBR Green Supermix (Bio-Rad Laboratories, CA, USA). Thermal cycling conditions were designed as follows: initial denaturation at 95°C for 3 min, followed by 45 cycles of 95°C for 10 s, and 60°C for 1 min. PCR cycle number at threshold is represented as Ct. Relative expression level of genes of interest were quantified by 2^−ΔΔCt^ method after normalization for GAPDH expressed in fold change as compared to corresponding uninfected control cells.

### 2.7. RT^2^ Profiler PCR Array

RT^2^ Profiler PCR Array Total RNA was analyzed using the human type I IFN response RT^2^ Profiler PCR array (catalog no. PAHS-016Z; QIAGEN), which profiles the expression of 84 gene transcripts that are known to be involved in the type I IFN response, as well as the expression of five housekeeping genes (ACTB, B2M, GAPDH, HPRT1, and RPLP0). In addition, one well contains a genomic DNA control, three wells contain reverse transcription controls, and three wells contain a positive PCR control. For each sample, 0.5 mg of RNA was reverse transcribed into cDNA using the RT^2^ first-strand kit (QIAGEN, UK). The cDNA was then mixed with the RT^2^ SYBR Green Master mix (QIAGEN, UK) and nuclease-free water. Next, 25 mL of the PCR mix was added to each well of the 96-well plate. All steps were done according to the manufacturer’s instructions. The RT-PCR reaction was run on an QuantStudio™ 3 Real-Time PCR System according to the following conditions: 95°C for 10 min, followed by 45 cycles of 95°C for 15 s and 60°C for 1 minute. Data analysis was conducted using a software-based tool (Applied Biosystems). Expression levels were quantified relative to the values obtained for housekeeping genes.

### 2.8. Flow cytometry

MCOs were washed 3 times in ice-cold DMEM/F12 and then dissociated by Papain. Briefly, MCOs were dissociated in a 1mL Papain for 45min mechanically disrupted every 15min. The cell suspension was then reverse filtered through a 100μm strainer (Thermo Fisher) and collected in a 15mL falcon tube. The Douncer was washed twice with 1mL DMEM/F12, added to the tube and the cells were centrifuged at 300g, at 4°C for 5min. Afterwards, cell counting was performed, and cells were resuspended in ice-cold FACs buffer (PBS supplemented with 2% BSA and 0.5mM EDTA). The cells were first blocked for 10min with Fc Blocker (cat#: BDB564765, BD Biosciences) and subsequently stained with antibodies (CD4-Pacific Blue cat#: 300521, Thermofisher; CCR5-APC cat#: 561748, BD Biosciences; CXCR4-PE/Cy7 cat#: 12G5, BioLegend; P2RY12-PE cat#: S16001E, BioLegend; GFAP-Alexa488 cat#: GA5, Thermofisher; MAP2-488 cat#: CL488-57015, Proteintech) or an isotype-matched antibody to determine the background fluorescence. After 30min at 4°C, the samples were acquired on Cytek Aurora (Cytek Biosciences), and the data analyzed using FlowJo v10.10 (Tree Star).

### 2.9. Phagocytosis assay

Primary human peripheral blood monocyte-derived macrophages (MDMs) and human iPSC-derived MCOs were incubated with pHrodo Red Zymosan BioParticles (5 µg/mL, cat #: P35364, Thermofisher) at 37°C for 4h or 24h, respectively. NCOs were then washed 2 times with warm medium to remove the unbound particles and re-suspended in colorless live image buffer. The uptake of particle conjugates was visualized using the Eclipse Ti2 microscope with a filter cube (Ex470/30, Em530/50) (Nikon camera).

### 2.10. ELISA

MCOs were plated in a 48-well plate (2 per well) and transfected with poly I:C. Culture supernatants were then collected and analyzed for IFNs and CC chemokines using a ELISA kits (DuoSet, R&D system). Briefly, 100μL of standards or samples from each well were incubated for 2h with detecting antibody prior to incubation with HRP conjugated 2^nd^ antibodies for 20min. The plates were read for the absorbance at 450nm with a wavelength correction at 540 nm was measured with a SpectraMax 340 plate reader (Molecular Devices, San Jose, CA, USA). The values were compared against a standard curve that was generated using known concentrations to calculate concentration in the samples.

### 2.11. Statistical analysis

GraphPad Prism 9.1 software (GraphPad Software, USA) was used to analyze data that were represented as means ± SD. The results were treated using two-way ANOVA, followed by multiple comparison tests for individual comparison to determine statistical significance.

## 3. Results and Discussion

To assess the structural developmental trajectory of the brain organoids, we monitored the sequential stages of brain organoid maturation, progressing from initial spherical structures to increasingly complex and dense formations (Figure 1A). Following embedding in Matrigel on day 30, brain organoids exhibited key developmental features, including the formation of a folded neuroepithelium-like surface, the presence of numerous rosette structures, and the establishment of a ventricular zone. By day 40, the cerebral organoids attained a diameter ranging from 3 to 5 mm, achieving macroscopic visibility. These morphological changes indicate successful differentiation and self-organization of neural progenitors, a hallmark of functional brain organoid development. MCOs’ size started stabilizing by day 50, with no further growth by day 60. Flow cytometry analysis showed that MCOs contained major brain cell types, including oligodendrocytes (37.5%), astrocytes (11.2%), neurons (42.9%) and microglia (2.5%) (Figure 1B). Importantly, the distributions of these brain cells in MCOs resemble the cellular diversity during human brain development. Furthermore, we observed that MCOs have phagocytosis function, being able to engulf pHrodo Red Zymosan particles (Supplemental Figure 1).

To study whether the MCOs are suitable for HIV-1 and innate immune response studies, we first examined the expression of TLRs and HIV-1 entry receptors (CD4, CCR5, and CXCR4) in MCOs. We observed that MCOs express all 10 TLRs (Supplemental Figure 2) and the HIV-1 entry receptors (Figure 1B). Building upon these findings, we next determined the impact of TLR3 activation on HIV-1 infection of MCOs. As shown in Figure 1C, while MCOs could be productively infected by live HIV-1_Bal_ strain, poly I:C treatment of MCOs potently suppressed HIV-1 infection. To study the cellular and molecular mechanisms for TLR3 activation-mediated HIV-1 inhibition in MCOs, we performed a comparative transcriptome analysis of control and poly I:C-treated MCOs. As demonstrated in Figure 2A, out of 84 genes in the IFN signaling pathway, 62 genes were significantly upregulated (>2 fold) by poly I:C treatment. Notably, there was about 10-fold increase of CD80 and CD86 gene expression, which are crucial in the initiation of antiviral immune responses. In addition, poly I:C treatment of MCOs resulted in the upregulation of gens of the cytokines (RANTES/CCL5, CXCL10, IL10, and IL15), viral detecting TLRs (TLR3, TLR7, TLR8, and TLR9), and IFNs/ISGs (IFN-β, IFN-λ, IFIT2, IFIT3, OAS1, OAS2, ISG15, MX1, and ISG20) (Figure 2A).

**Figure 2.**
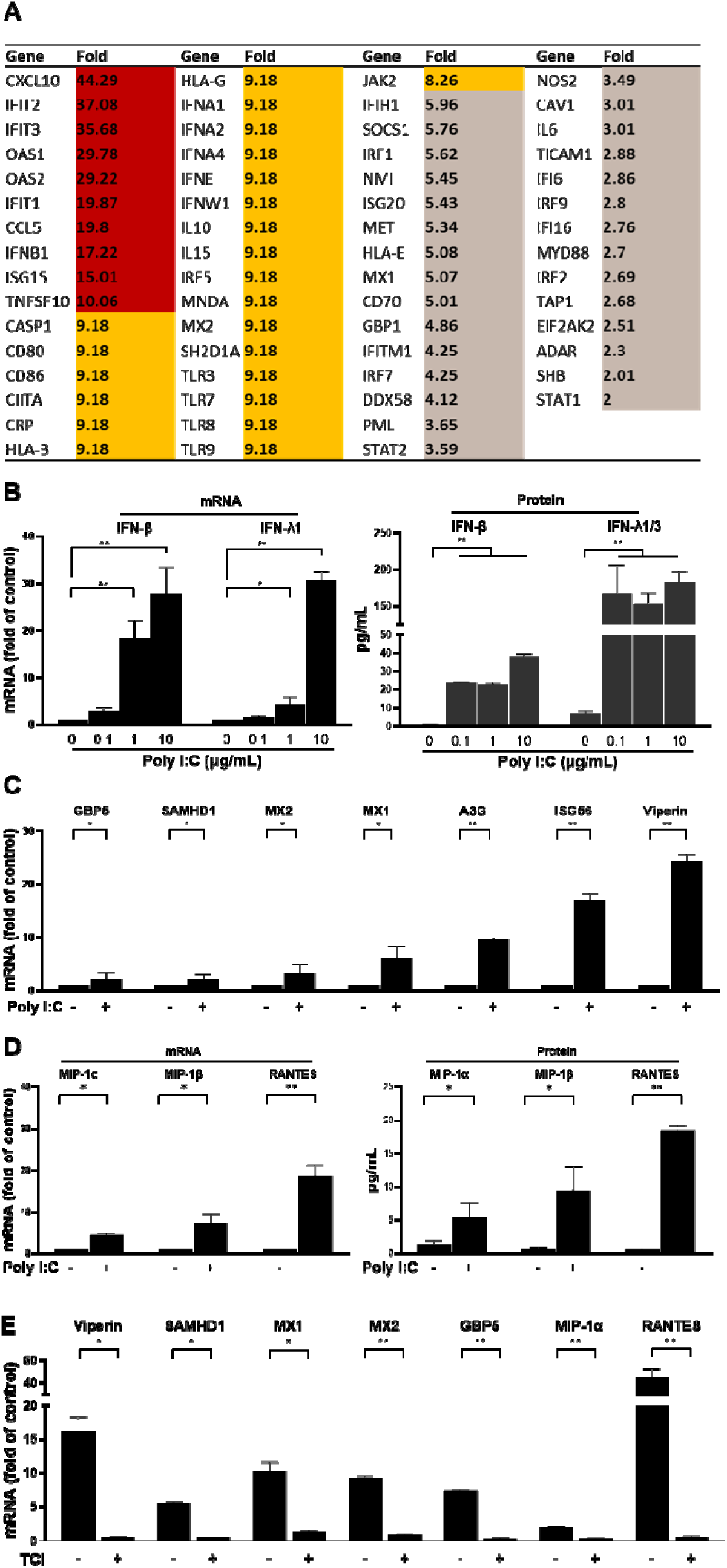
TLR3 activation of MCOs upregulates type I IFN pathway genes and induces CC chemokines. (**A**) Poly I:C treatment upregulates type I IFN pathway genes in MCOs. MCOs were treated with poly I:C (10 µg/mL) for 24h and mRNA levels of 84 genes were assessed with a Type I IFN RT² Profiler PCR array. The data shows gene expressions as absolute fold changes, with different colors indicating upregulation levels: brown (< 6-fold), orange (8 -10 fold), and red (> 10-fold). (**B**) TLR3 activation of MCOs induces IFN-β and IFN-λ1. MCOs were treated with poly I:C at the indicated doses, and the supernatants or cell lysate were collected for RT-PCR and DuoSet ELISA at 24h post-treatment. Data shown are the mean ± SD of triplicate cultures, representative of four independent experiments (*P < 0.05, **P < 0.01). (**C**) TLR3 activation of MCOs induces ISGs. MCOs were treated with poly I:C (10 µg/ml), and the induction of the indicated ISGs was measured by RT-PCR at 24h post-treatment. Data is presented as the fold change relative to the untreated MCOs. Data shown are the mean ± SD of triplicate cultures, representative of four independent experiments (*P< 0.05, **P< 0.01). (**D**) TLR3 activation of MCOs induces CC chemokines. MCOs were treated with poly I:C (10 µg/mL) and CC chemokines were analyzed by RT-PCR and ELISA at 24h post-treatment. Data shown are the mean ± SD of triplicate cultures, representative of four independent experiments (*P< 0.05, **P< 0.01). (**E**) TLR3 inhibitor blocks poly I:C-mediated induction of the ISGs and CC chemokines. MCOs were pretreated with or without TLR3/dsRNA complex inhibitor (TCI, 12.5µM) for 4h prior to addition of poly I:C (10µg/ml) to the cultures. The mRNA expression of the antiviral ISGs and chemokines were analyzed by RT-PCR. Data shown are the mean ± SD of triplicate cultures, representative of two independent experiments (*P < 0.05, **P < 0.01).

Our earlier studies using human iPSC-derived microglia showed that TLR3 activation could induce the intracellular antiviral state characterized by the upregulation of IFNs, antiviral ISGs, and the CC chemokines which can restrict HIV-1 replication (Wang et al., 2023; Yu Zhou et al., 2010). To further study the immune response of MCOs to TLR3 activation by poly I:C, we investigated whether TLR3 activation of MCOs could induce IFNs, antiviral ISGs and CC chemokines. We observed that treatment of MCOs with poly I:C resulted in a dose-dependent increase of IFN-β and IFN-λ at both mRNA and protein levels (Figure 2B). This selective induction of IFN-β and IFN-λ may be attributed to distinct signaling pathways or cell-type specific responses whin MCOs. We also showed that poly I:C -treated MCOs expressed higher levels of antiviral ISGs (MX2, GBP5, MX1, SAMHD1, Viperin, and ISG56) than untreated MCOs (Figure 2C). In addition, TLR3 activation of MCOs elicited production of CC chemokines (MIP-1α, MIP-1β, and RANTES) (Figure 2D), the ligands of HIV-1 entry coreceptor (CCR5). To confirm that TLR3 is responsible for poly I:C- mediated IFN signaling pathway activation in MCOs, we treated MCOs with a TLR3 inhibitor (TLR3/dsRNA complex inhibitor, TCI) at non-cytotoxicity concentration (Supplementary Figure 3) prior to poly I:C stimulation. As demonstrated in Figure 2E, TCI effectively blocked TLR activation-mediated induction of IFN-β/IFN-λ, antiviral ISGs (ISG15, MX2, OAS1, GBP5, and Viperin), and CC chemokines (MIP-1α, MIP-1β, and RANTES) in MCOs.

In summary, the observations presented in this paper provide experimental evidence that MCOs possess functional TLR3, stimulation of which by poly I:C activates IFN signaling pathway and induces the production of multiple antiviral cellular factors against HIV-1. These findings are clinically relevant and significant, as they demonstrate that iPSC-derived cerebral organoids integrated with microglia are a valuable model not only for investigating human brain innate immunity against HIV-1 infection, but also for exploring TLR3 as a potential HIV-1 therapeutic target in the brain.

## CRediT authorship contribution statement

Qian-hao Xiao: Methodology, Formal analysis, Investigation, Writing – original draft, Writing – review & editing. Xu Wang: Investigation, Writing – review & editing, Supervision. Priyanka Sarkar: Methodology, Formal analysis. Xiao-Long Wang: Methodology, Formal analysis. Li Song: Methodology, Formal analysis. Binhua Ling: writing-review and editing. Wen-Hui Hu: Investigation, Resources, Writing – review & editing, Funding acquisition. Wen-Zhe Ho: Methodology, Conceptualization, Formal analysis, Resources, Writing – review & editing, Project administration, Supervision, Funding acquisition.

## Acknowledgments

The authors declare that this study received funding from NIH (R01 DA051893, R01 DA058536 and R01 MH134402). The funder was not involved in the study design, collection, analysis, and interpretation of data, writing of this article or the decision to submit it for publication.

## Data availability statement

The data that support the findings of this study are available from the corresponding author upon reasonable request.

