## Supplemental information for "TLR3 Activation of Microglia-Containing Cerebral Organoid Induces Antiviral Factors against HIV-1 Infection"

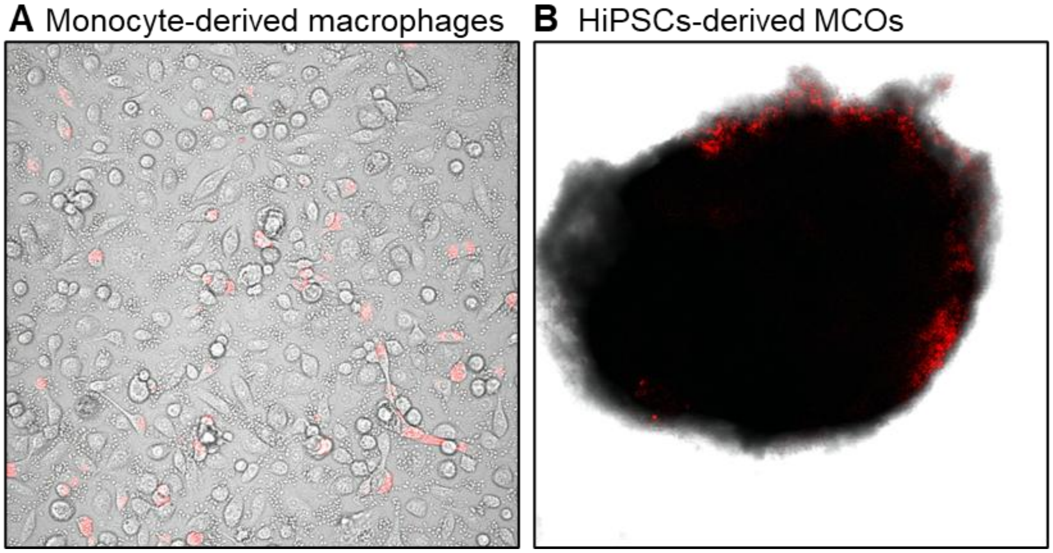


**Supplementary Figure 1. MCOs showed phagocytic properties as macrophages.** Phagocytosis capacity was analyzed in (**A**) primary human peripheral blood monocyte-derived macrophages (MDMs) and (**B**) MCOs (day 50). MDMs and MCOs were incubated with 5 µg/mL pH-sensitive Zymosan for 4h and 24h, respectively, and imaged by a Nikon confocal microscopy.

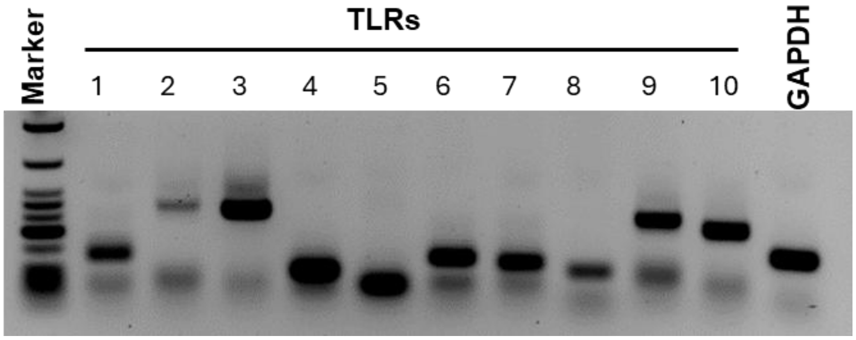


**Supplementary Figure 2. Detection of TLRs (1-10) mRNA by RT-PCR in MCOs (day 50).** cDNAs (amplified for 35 cycles) were separated on a 3% agarose gel containing ethidium bromide. 1-kb DNA ladder standard was used as a size marker.

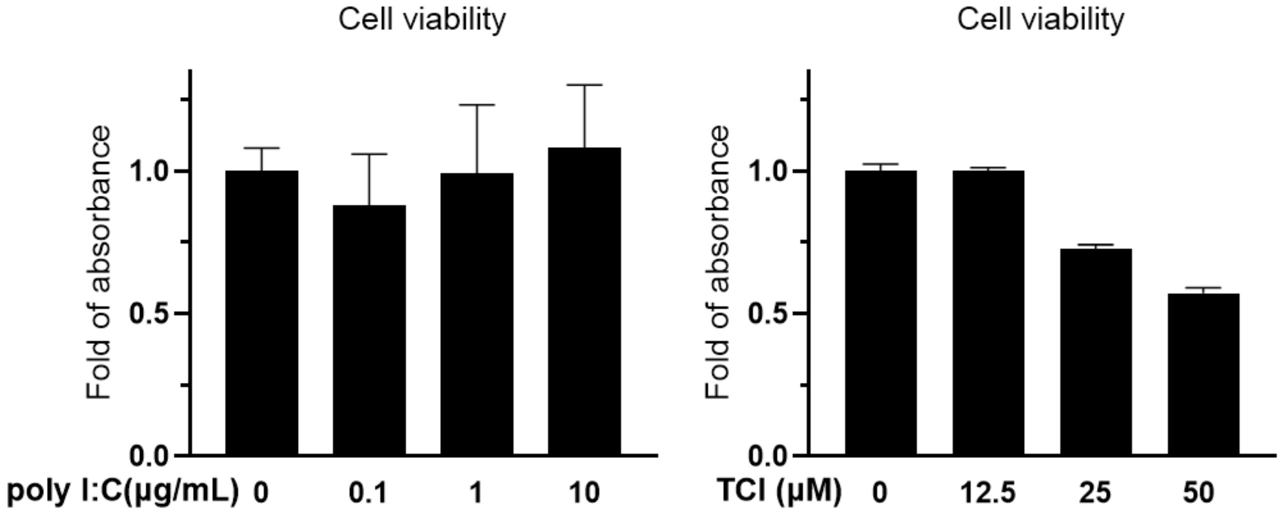


**Supplementary Figure 3. Effect of polyI:C or TLR3/dsRNA complex inhibitor (TCI) on cell viability of MCOs.** MCOs (day 50) were dissociated into single cells and treated with poly I:C or TCI at indicated concentrations for 48h. Cell death was quantified by cell proliferation assay Kit. Data shown are the mean ± SD of triplicate cultures and representative of two independent experiments.
